# Identification of Disaggregated Hotspots of Child Morbidity in Bangladesh: An Application of Small Area Estimation Method

**DOI:** 10.1101/699538

**Authors:** Sumonkanti Das, Bappi Kumar, Luthful Alahi Kawsar

## Abstract

Acute respiratory infection (ARI) and diarrhoea are two major causes of child morbidity and mortality in Bangladesh. National and regional level prevalence of ARI and diarrhoea are calculated from nationwide surveys; however, prevalence at micro-level administrative units (say, district and sub-district) is not possible due to lack of sufficient data. In such case, small area estimation (SAE) methods can be applied by combining a survey data with a census data. Using a SAE method for dichotomous response variable, this study aims to estimate the proportions of under-5 children experienced with ARI and diarrhoea separately as well as either ARI or diarrhoea within a period of two-week preceding the survey. The ARI and diarrhoea information extracted from Bangladesh Demographic and Health Survey 2011 are used to develop a random effect logistic model for each of the indicators, and then the prevalence is estimated adapting the World Bank SAE approach for the dichotomous response variable using the 5% data of the Census 2011. The estimated prevalence of each indicator significantly varied by district and sub-district (1.4-11.3% for diarrhoea, 2.2-11.8% for ARI and 4.3-16.5% for ARI/diarrhoea at sub-district level). In a number of districts and sub-district, the proportions are found double the national level. District and sub-district levels spatial distributions of the indicators might help the policy makers to identify the vulnerable disaggregated and remote hotspots. Particularly, aid industries can provide effective interventions at the highly vulnerable spots to overcome the gaps between micro and macro level administrative units.

## Introduction

Diarrhoea and acute respiratory infection (ARI) are recognized as important causes of global child morbidity and mortality. Pneumonia (a form of ARI) and diarrhoea remain major causes of child death. The impact of these two diseases is about 29% of all under-5 child deaths causes loss of 2 million young lives each year [1]. Pneumonia alone accounts for about 16% of these young child deaths. The number of deaths due to pneumonia can be reduced by the early diagnosis and treatment of ARI. The integrated Global Action Plan for Pneumonia and Diarrhoea (GAPPD) aims to reduce mortality from pneumonia and diarrhoea among under-5 children to fewer than 3 per 1000 and 1 per 1000 live births respectively by 2025 [2]. The United Nation (UN) has set a target under the third sustainable development goal (SDG) to end the epidemics of water-borne diseases and other communicable diseases by 2030 (target 3.3) with the aim of achieving the SDG target of ending preventable deaths of under-5 children to reduce child mortality to below 25 per 1,000 live births (target 3.2). Recently WHO reported that under-5 child mortality rates reduced to 41 per 1000 live births in 2016 from 93 per 1000 live births in 1990, however still now everyday on an average 15,000 children die before reaching their fifth birthday [3]. About half of these under-5 child deaths can be prevented through simple and cost-effective interventions at the proper time.

As per GAPPD, the solutions to tackling pneumonia and diarrhoea do not require major advances in technology since proven interventions already exist. Children are dying because the provided services are not enough to meet the demand and children at higher risk are not being reached. Use of effective interventions such as exclusive breastfeeding for the first 6 months, proper life-saving treatment for the children with suspected pneumonia and providing oral rehydration therapy to the children with diarrhoea are sufficient to achieve these goals [2]. Thus, identifying the children at greatest risk, hardest to reach and most neglected, and targeting them with interventions of proven efficacy will enable us to close the gap, ultimately ending the heavy toll of preventable child deaths.

The prevalence of ARI and diarrhoea at national and divisional levels are usually estimated from a nationwide household survey. In Bangladesh, such nationwide data on ARI and diarrhoea information are collected through household surveys conducted by Bangladesh Bureau of Statistics (BBS), and through the Demographic and Health Survey (DHS). Diarrhoea and ARI data are collected by asking mothers whether their children experienced a diarrhoea episode or ARI symptom during a two-week period preceding the survey. According to the seven Bangladesh DHS (BDHS) surveys during the period of 1993-2014, the prevalence rates of diarrhoea and ARI have been improved in Bangladesh over the period of 1990-2010, however, there were no steady declining trend. The prevalence of diarrhoea was around 20% over the period of 1993-2004 with a decline only in 1996 (about 13%), however the rate was found stable at around 5% in the later surveys. For ARI, there was a somewhat steady decline in the prevalence over the period except in 2007, when the rate was unexpectedly jumped to about 10% [4]. In the last 2014 BDHS, the episodes of ARI and diarrhoea were found around 5% for both cases [4], while the rates were 6% and 5% respectively in 2011 BDHS [5]. The distribution of diarrhoea and ARI prevalence at division level does not show any specific declining trends [4-10]. Collection of survey data at different time-periods may be one of the reasons, which suggest some seasonal effects in the prevalence. An approximate declining trend in both diarrhoea and ARI prevalence are observed only for *Rajshahi* (and also *Rangpur*) division during the 1993-2014 period. Interestingly, water-prone divisions *Chittagong*, *Barisal* and *Sylhet* were most likely vulnerable to diarrhoea and ARI diseases in most of the surveys. Although division level estimates of diarrhoea and ARI prevalence are estimable from the survey data, those at disaggregated levels (such as district and sub-district) are not estimable solely from the survey data due to the limited number of observations at the desired micro level. Consequently, it is impossible for the government to find the disaggregated hotspots highly vulnerable to ARI and diarrhoea. Identifying such hotspots might help the aid industries concerned targeting efficient interventions.

A few district level studies on diarrhoea prevalence are found in literature. In a cross-sectional study covering seven vulnerable districts of Bangladesh which are prone to cyclone, flood, and salinity (*Bagerhat*, *Barguna*, *Cox’s Bazar*, *Faridpur*, *Khulna*, *Satkhira*, and *Sirajganj*), 10.3% under-5 children (95% CI 9.16-11.66) experienced with diarrhoea in 2012 during the preceding month of interview [11]. Though they have district-specific data, they did not reported district specific estimates of diarrhoea prevalence. No study has also been found conducted at the sub-district or lower administrative units. Most of the studies of ICDDR’B are related to environmental and clinical risk factors of different types of diarrhoeal diseases [12, 13]. Using BDHS data, there are studies on modelling diarrhoea or ARI prevalence based on logistic or logistic mixed model for determining their risk factors [14, 15], not for prediction purpose since information of the risk factors are not available in the census data.

Since districts and sub-districts are ignored in the sampling design for the nationally representative household survey, the estimates of diarrhoea and ARI prevalence or any other target parameters are not estimable at these levels. The BDHS 2011 data covers all the 64 districts but only 396 out of 544 sub-districts. Since the sample sizes both at district and sub-district levels are too small to have efficient estimates, the design-based direct estimates are not reasonable to use [16]. Consequently, the policy makers are unable to do their plan focusing on the district or its lower administrative hierarchies.

Small area estimation (SAE) is a statistical technique to obtain estimates of a target parameter with better precision for disaggregated administrative units of a country. The basic idea of SAE method is to model survey data statistically [16]. Such modelling may include use of a recent census or administrative data. Survey data consist of the target variable and a regression model is specified with some explanatory variables which are common in both survey and census data. The SAE methods are broadly two types based on the availability of the explanatory information. If unit level explanatory variables (such as children level variables used in this study) are available for all the population units (or a sample of the census), unit-level SAE methods use unit-level models such as nested error regression model [17]. While if area-level aggregate statistics extracted from census or any administrative data source are available, area-level SAE methods apply area-level models such as Fay-Herriot model [18]. One of the major problems of area-level method is that the estimated standard errors are assumed known, though there are always significant number of small areas (say here sub-districts) with unreliable standard errors since they are calculated based on small sample sizes. Also for some small areas, the estimates as well as standard errors are not available due to zero observation. Another problem is that the area-level model SAE method can be used for estimating the prevalence only for that level. Higher-level estimates such as division level can be estimated by aggregating the district level estimates through a suitable weighted method, however not possible to estimate the prevalence at lower administrative units like sub-district. The unit-level SAE method has advantages for avoiding the two mentioned problems and so a unit level SAE approach has been followed to estimate the prevalence of the considered health indicators in this study.

The World Bank has been utilising a unit-level SAE method known as ELL after the authors Elbers, Lanjouw, and Lanjouw [19] for poverty and nutrition mapping in many developing countries including Bangladesh. The basic idea of the ELL methodology is to develop a regression model using a continuous response variable (such as per capita consumption expenditure, weight-for-age Z-score) from survey data and apply it to a census or administrative data source. Since the variable of interest for diarrhoea and ARI prevalence is dichotomous (whether a child has experienced with diarrhoea or not during a period) instead of continuous, the ELL methodology cannot be implemented without modification. However, the basic idea can be implemented after developing a generalized linear mixed model (GLMM) more specifically a random effect logistic model for the dichotomous response variable [20]. The main difficulty is to develop a proper GLMM model incorporating the available survey data with a recent census or administrative data for a country. Another problem arises in developing a proper logistic model when the event is less frequent among the study population [21]. In this study, developing a proper (mixed effect) logistic model with sufficient explanatory variables might be problematic since occurrence of diarrhoea (and also for ARI) was found only for about 5% children in the BDHS 2011 data.

Recently Das, Chandra and Saha [22] conducted a SAE study on district level diarrhoea prevalence in Bangladesh employing the area-level SAE method. District specific direct estimates of diarrhoea prevalence with their standard errors were calculated based on the design-based direct estimator using the data of BDHS 2014. These direct estimates were used as the response variable and a number of district-specific variables collected from the Census 2011 reports are used as the explanatory variables. Though division level diarrhoea prevalence can be calculated from these district-level estimates, it is not possible to estimate sub-district specific diarrhoea prevalence. For estimating sub-district level diarrhoea prevalence, sub-district (or more lower administrative level) specific Fay-Herriot model is required to develop that might be difficult due to lack of efficient direct estimates at that level (again the issue of very small number of under-5 children at sub-district level in the survey data). Thus, the main aim of this study is to estimate the prevalence of ARI and diarrhoea for under-5 children at district and sub-district levels using a unit-level SAE technique for dichotomous response variable. In addition, the aims include estimating the proportion of children suffering from at least one of the two indicators during the 2-week period (hereafter refereed as ARI/diarrhoea) at disaggregated levels. Finally, disaggregated level spatial distributions of all the three indicators are mapped to highlight the most vulnerable hotspots. The rest of the paper is set up as follows: Section 2 describes survey and census data used in this study; Section 3 describes a design-based direct estimator and a model-based SAE estimator for binary response variable based on the ELL method; Section 4 illustrates the fitted models, spatial distribution of the prevalence, and explores the characteristics of the considered SAE estimator; and concluding remarks are given in Section 5.

## Data description

Diarrhoea and ARI data used in this study are extracted from the 2011 BDHS survey. The main reasons for using BDHS 2011 (instead of recent BDHS 2014) is it’s concurrency with the Census 2011 and also the hierarchy of the administrative units are available down to sub-district (sub-districts are unidentifiable in BDHS 2014). The survey data is collected following a two-stage stratified sampling design by covering all 7 divisions, 64 districts, and 396 (out of 544) sub-districts. In the BDHS 2011, a total of 8341 children were found whose ARI and diarrhoea information were available [5].

Full census data of Bangladesh is unavailable for academic purposes; however, 5% of the full census data is available from BBS. Socio-demographic characteristics such as age, sex, education, schooling, employment, disability and housing characteristics such as house type, source of drinking water, sanitation are available in the census data. A number of contextual variables at district and sub-district levels can be created using the individual and household level data of the 5% census data. In the model specification, these contextual variables are used in addition to those variables at child level common in both survey and census. The contextual variables used in the model development for capturing the variation at the district and sub-district levels are shown in Appendix Tables 1 and 2. In model development, two-way interactions of residence, division, sex, and age are utilized to develop the best models. The interaction of division and residence also covers the sampling design of the BDSH.

**Table 1.**
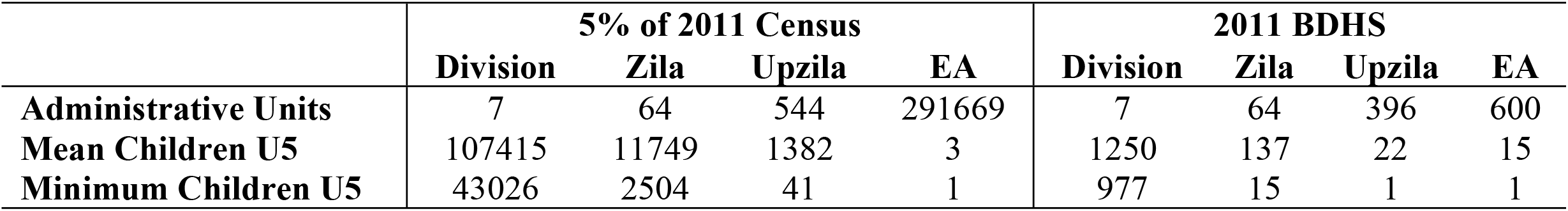
Structure of administrative units and children in the Census 2011 and BDHS 2011.

**Table 2.** Summary statistics and diagnostics of the fitted logistic (GLM) and random intercept logistic (GLMM) models for Diarrhoea, ARI, and ARI/diarrhoea prevalence, BDHS 2011 ♀.

| Indicators | n | P | Model | AIC | logLik | $\sigma_u^2$ | LRT of $H_0: \sigma_u^2 = 0$ | AUC (%) |
| --- | --- | --- | --- | --- | --- | --- | --- | --- |
| Diarrhoea | 8341 | 21 | GLM | 3111.8 | -1534.9 | - | $\chi^2$: 3.7120 | 54.86 |
|  |  |  | GLMM | 3110.1 | -1533.0 | 0.1687 | p-value: 0.0215 | 71.55 |
| ARI | 8341 | 22 | GLM | 3680.2 | -1818.1 | - | $\chi^2$: 6.2343 | 61.11 |
|  |  |  | GLMM | 3676.0 | -1815.0 | 0.1921 | p-value: 0.0063 | 72.26 |
| ARI/Diarrhoea | 8341 | 21 | GLM | 5371.6 | -2664.8 | - | $\chi^2$: 2.7519 | 58.41 |
|  |  |  | GLMM | 5370.8 | -2663.4 | 0.0791 | p-value: 0.0448 | 65.78 |

♀ n=sample size, p= # of covariates, $\sigma_u^2$ =cluster-level variance component, LRT= likelihood ratio test, AUC= Area under the receiving operating characteristics curve

The structure of administraive units and the distribution of children in BDHS 2011 and Census 2011 are shown in Table 1. The mean and minimum number of children at district (137 and 15) and sub-district (22 and 1) levels indicates that realiable estimates are not possible from the BDHS survey at these disaggrgated levels, particulalrly at the sub-district level. Since the BDHS data is collected through cluster sampling design and alos cluster-specific multilvel models will be developed, cluster specific information are also examined. Table 1 shows that mean number of children at the cluster level in the survey is higher than that in the census data. As per definition of enumeration area (EA) which is the cluster in the survey, an EA/cluster consits of on average 110-120 households. In the survey 30 households were covered, while 5% of these households are selected. As a consequence, the mean number of under-5 children is very low in the census part.

In model development part, the number of children experienced with diarrhoea is found zero for about three-fifths clusters (351 out of 600), for ARI and ARI/diarrhoea the proportios are respectively 50.1% and 32.3% (301 and 194 out of 600). These practical issues would be problemtic for developing an appropriate multilevel logistic model. This issue is discussed in model development section.

## Statistical Methodology

Let the occurrence of an event for *k*^*th*^ child belonging to *j*^*th*^ cluster of *i*^*th*^ area is denoted by *y*_*ijk*_, which takes value 1 if the child experienced the event (say diarrhoea) preceding the last two-week of the survey date and 0 if the child did not experience the event. The target is to calculate proportion of under-5 children who experienced a target event during the period. The proportion is aimed to calculate at three aggregation levels division, district and sub-district in Bangladesh. The survey data is representative at the division level and so design-based direct estimator provide unbiased and consistent division level estimates but the problem arises for the other two disaggregated levels. The SAE method has been applied for estimating proportions at all the three levels. The design-based direct estimator and a SAE estimator for binary response variable based on unit-level SAE method have been briefly explained in the following two sub-sections.

### Design-based Direct Estimator

The Horvitz-Thompson estimator for estimating mean and total of a super-population in a stratified sample are utilized to obtain design-based direct estimates of target parameter for small areas using the response values available in the survey data. The design-based direct estimator (denoted by DIR) for the target proportion *P_i_* for area *i* is

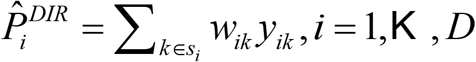

where 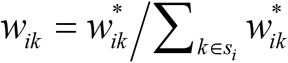 is normalized survey weights for *k*^*th*^ child belonging *i*^*th*^ district with 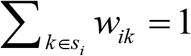 and 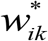 is the survey weight (inverse of the inclusion probability). Following Särndal *et al.* [23], the design-based variance of the direct estimator 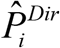 can be approximated by,

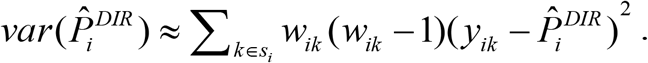

This direct survey estimator is design-unbiased but based on the non-representative area-specific sample data. Consequently, the direct estimator becomes unreliable due to area specific small sample size and also for some areas with no sample data. In BDHS 2011 data, though all the districts are covered, sample size for a significant number of districts are very small since only a few clusters are covered for those districts. Also for some districts, the number of children experienced with an event (say, diarrhoea) is found zero, which provides zero prevalence and zero standard error. The model-based SAE methods that ‘borrow strength’ via statistical models overcome these practical issues for calculating reliable small area estimates [24].

### Model-based SAE Estimator

Suppose 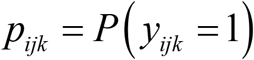 represents the probability of having diarrhoea for *k*^*th*^ child belonging to *j*^*th*^ cluster of *i*^*th*^ area. The first target is to develop a nested-error logistic regression model which is a special case of generalized linear mixed model (GLMM) as

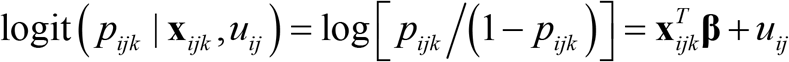

Where **x**_*ijk*_ is vector of explanatory values, **β** is vector of regression parameters, and *u*_*ij*_ corresponds to cluster specific random errors respectively. The random errors are usually assumed to be independent and identically distributed with mean zero and constant variance 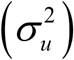. The regression parameters **β** and variance component 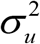 can be estimated restricted maximum likelihood method (REML). The regression model can be extended to higher level (such as area-specific effect here), however the ELL methodology assumes heterogeneity at cluster-level rather than target area levels [19]. For estimating the target parameters with their root mean squared error (RMSE), the estimated regression coefficients, variance components, residuals, and the explanatory information for all the under-5 census children are used as input in a parametric or non-parametric bootstrap procedure of the ELL method. The basic steps of the parametric or non-parametric bootstrap procedure are briefly explained following Das and Haslett [25].
Step 1: fit random effect logistic model to obtain regression coefficients 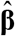 with their estimated variance-covariance matrix 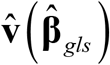, and cluster-specific residuals with the variance component 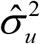.
Step 2: generate regression parameters **β**^*^ from a suitable sampling distribution, say the multivariate normal distribution 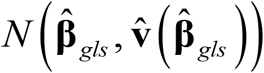;
Step 3: generate cluster-specific random errors (except the children level) from a suitable parametric distribution such as 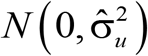 or t-distribution. In case of non-parametric bootstrap procedure, draw cluster-specific random errors by resampling via simple random sampling with replacement (SRSWR) from their empirical distribution (i.e., from the estimated cluster-specific sample residuals stored in Step 1);
Step 4: generate bootstrap response values 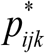 using the generated regression parameters and the cluster-specific random errors. The generated response values 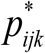 are then aggregated to estimate the area-specific parameter of interest say 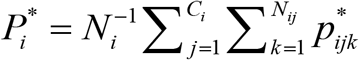 where *N_i_* and *N_ij_* are area and cluster specific number of under-5 census children.

The steps 2-4 are iterated for a large number of times say *B*=500 and then the mean and variance of these *B* estimates are considered as the final estimates and their RMSEs respectively as

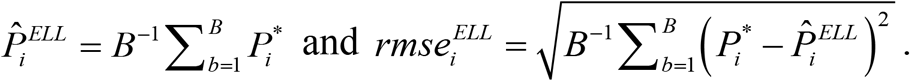

This estimator is denoted by ELL in the text since the estimator is developed based on the ELL approach but used for binary response variable. In the original development of ELL method for continuous response variable, heteroscedasticity has been considered at the unit-level (say children here). If the response variable is weight-for-age Z-score, this heteroscedasticity issue can be employed. Since, the response variable is binary, the heteroscedasticity at the unit-level is not considered here.

## Results and Discussion

Fixed effect logistic models (GLM) and random intercept logistic models (GLMM) are developed using children demographic characteristics, household characteristics, place of residence, regional settings, and a number of contextual variables. The final selected models for these three health indicators are shown in Appendix Table 1. The final models with their corresponding inputs are utilized in the ELL approach to estimate the proportion of each health indictor with their RMSEs and coefficients of variation (CVs). Table 2 shows the comparison of GLM and two-level GLMM models, which indicates that two-level GLMM models are performing better than the fixed effect GLM model in terms of AIC, Likelihood Ratio Test (LRT), and area under the ROC curve (AUC) for each of the three child health indicators. The AUC values indicate that the GLMM can classify children health status more correctly than the GLM in all cases; particularly for ARI about 73% children are correctly classified.

Although the cluster-specific random errors are assumed to follow the normal distribution, the normality assumption is not satisfied for diarrhoea and ARI separately, however, the normality assumption is approximately satisfied for ARI/diarrhoea (see Q-Q plots in Fig 1). The distribution of clusters with zero prevalence (58.1% and 50.2% respectively for diarrhoea and ARI) may be one of the reason for such non-normal distribution of residuals obtained from the models of diarrhoea and ARI prevalence. Although the residual distributions are found non-normal, their variances are found approximately homogeneous (see distributions of residuals in Fig 1). To avoid the impact of non-normal residuals, non-parametric bootstrap procedure assuming constant variance of residuals are employed in the prediction of these health indicators.

**Fig 1:**
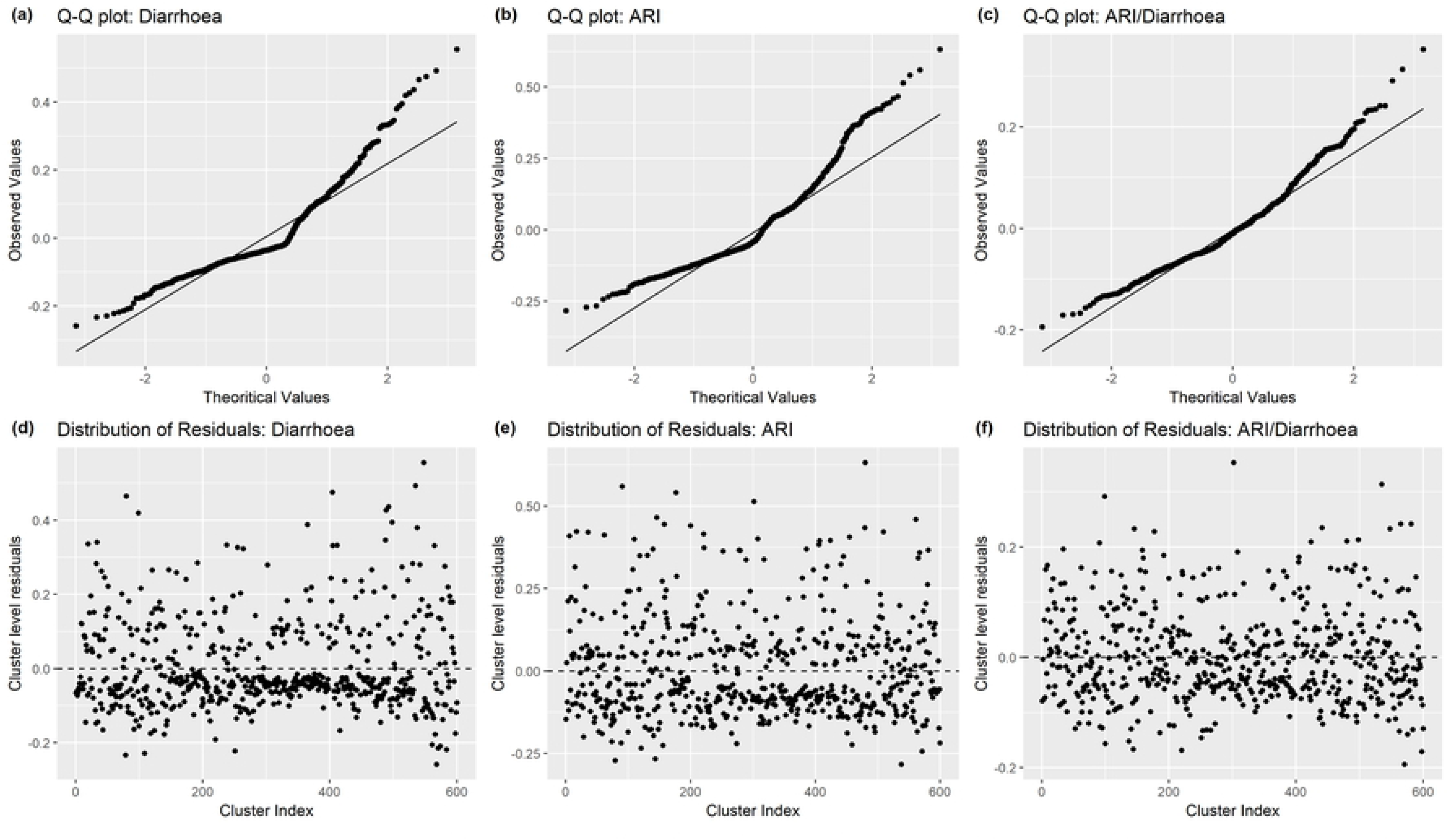
Normal Q-Q plots and distributions of the cluster-specific residuals for the fitted models of diarrhoea, ARI and ARI/diarrhoea

To examine the performance of SAE method at the higher administrative levels, division level estimates are also estimated and compared with the estimates calculated by the design-based DIR estimator. The bar plots with 95% confidence interval (CI) shown in Fig 2 (plots (a), (c) and (e)) indicate that the ELL estimator provides very similar estimates as the DIR estimator at division level, however the ELL estimator shows higher accuracy measured by the coefficient of variation (CV, %). The bar plots of CVs by division in Fig 2 (plots (b), (d) and (f)) show that the ELL estimator provides considerably lower CVs than the DIR estimator for most of the divisions except *Dhaka* and *Chittagong* division, where the sample number of children are considerably higher than the other divisions. The 95% CI lines also show that the DIR estimator provides higher confidence interval for most of the division due to producing higher standard errors.

**Fig 2:**
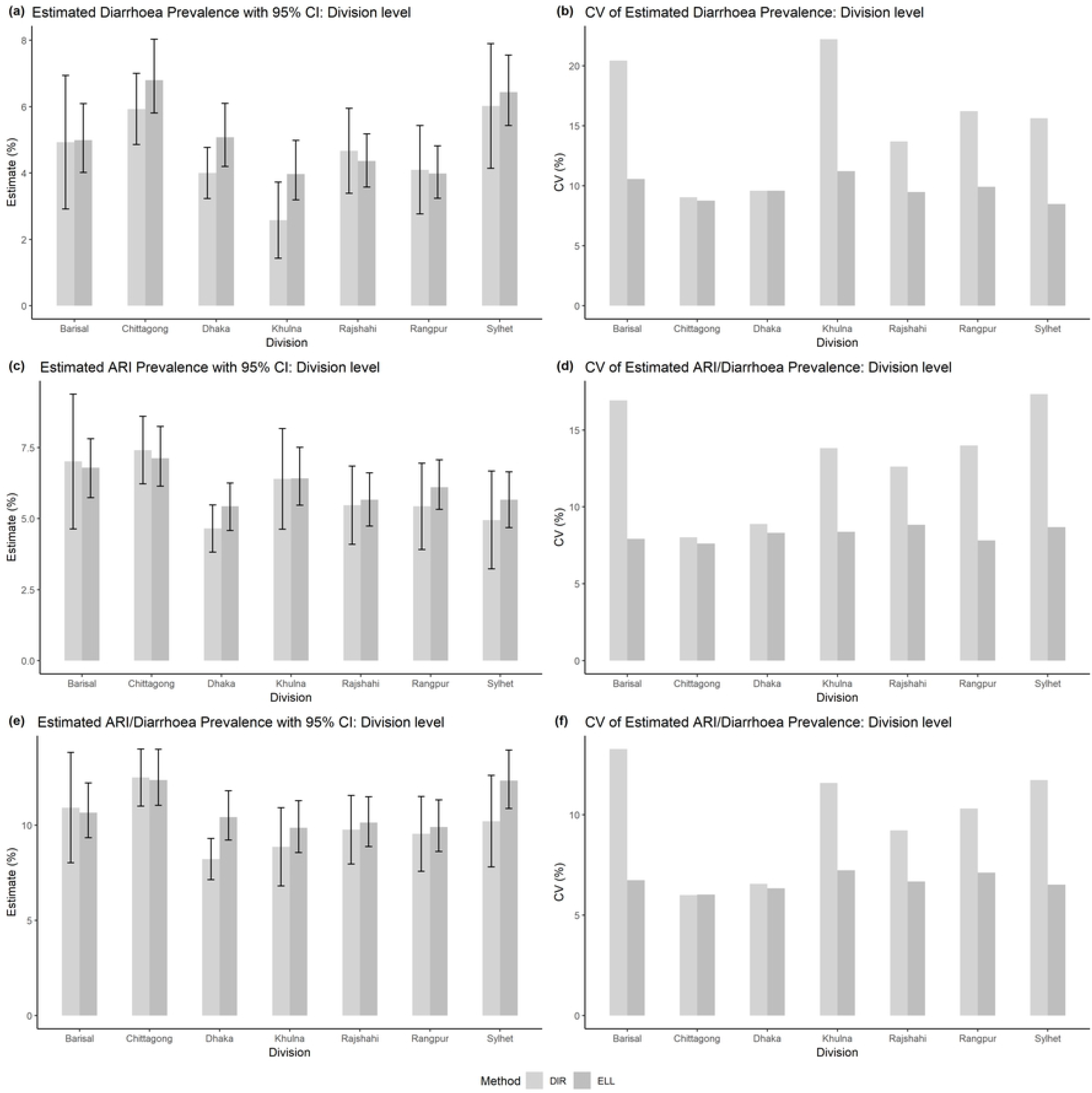
Prevalence of diarrhoea, ARI, and ARI/diarrhoea (plots (a), (c), (e) respectively) among under-5 children at division level in Bangladesh with their 95% CI and also their coefficient of variations (CV) (plots (b), (d), (f) respectively) estimated by direct (light grey bars) and ELL (grey bars) estimators

The division level prevalence of diarrhoea, ARI, and ARI/diarrhoea estimated by the ELL estimators indicate that the children living in *Chittagong* division are highly vulnerable to all the three indicators (about 6.5%, 7.5% and 12% respectively), followed by those living in *Sylhet* and *Barisal* divisions. On the other hand, the children of *Khulna* division had lower prevalence of diarrhoea (near about 4%) but they were more affected by ARI (near about 6%). The bar plot correspond to ARI/diarrhoea indicate that near about 10% children living in *Dhaka*, *Khulna*, *Rajshahi*, and *Rangpur* divisions had experienced with either ARI or diarrhoea, though there were some difference in terms of diarrhoea and ARI prevalence such as lowest diarrhoea prevalence in *Khulna* and lowest ARI prevalence in *Dhaka*.

The district level prevalence of diarrhoea, ARI and ARI/diarrhoea estimated by the ELL estimator are plotted against those prevalence estimated by the DIR estimator for assessing the unbiasedness of the ELL estimator. The bias diagnostic plots with the y=x lines and regression lines shown in Fig 3 (plot (a) for diarrhoea, (c) for ARI and (e) for ARI/diarrhoea) indicate that the ELL estimator provides approximately unbiased estimates compared to the DIR estimates. The bias diagnostic plots may indicate that the ELL estimator did more shrinkage to the average estimates than the DIR estimator did. The main reason might be very high prevalence estimated by the DIR estimator for some districts with small number of children and also for some districts zero prevalence (for diarrhoea and ARI). For comparing the accuracy of the district level prevalence estimated by the ELL and DIR estimators, the CVs of estimated diarrhoea, ARI and ARI/diarrhoea prevalence are plotted against the district index ordered by the number of children in Fig 3 (plots (b), (d), and (f) respectively). The plots for CVs indicate that the ELL estimator provides considerably lower CVs than those of DIR estimator as expected. It is noted that for the districts with zero prevalence, standard error and CV are not possible to calculate and so there are some discontinuities in the CV lines of DIR estimator for diarrhoea and ARI prevalence. This is another advantage of the SAE technique, which provides estimates with accuracy for those areas having no information in the sample data. The summary statistics of the district level prevalence indicate the diarrhoea prevalence varied within 2.69-5.69%, ARI prevalence within 4.26-8.92%, and ARI/diarrhoea 7.71-13.73% (please see Table 3). The maximum CVs for the district level prevalence are estimated as respectively 20.81%, 18.25%, and 12.10%, which indicate that the accuracies of the estimated prevalence are very good.

**Fig 3:**
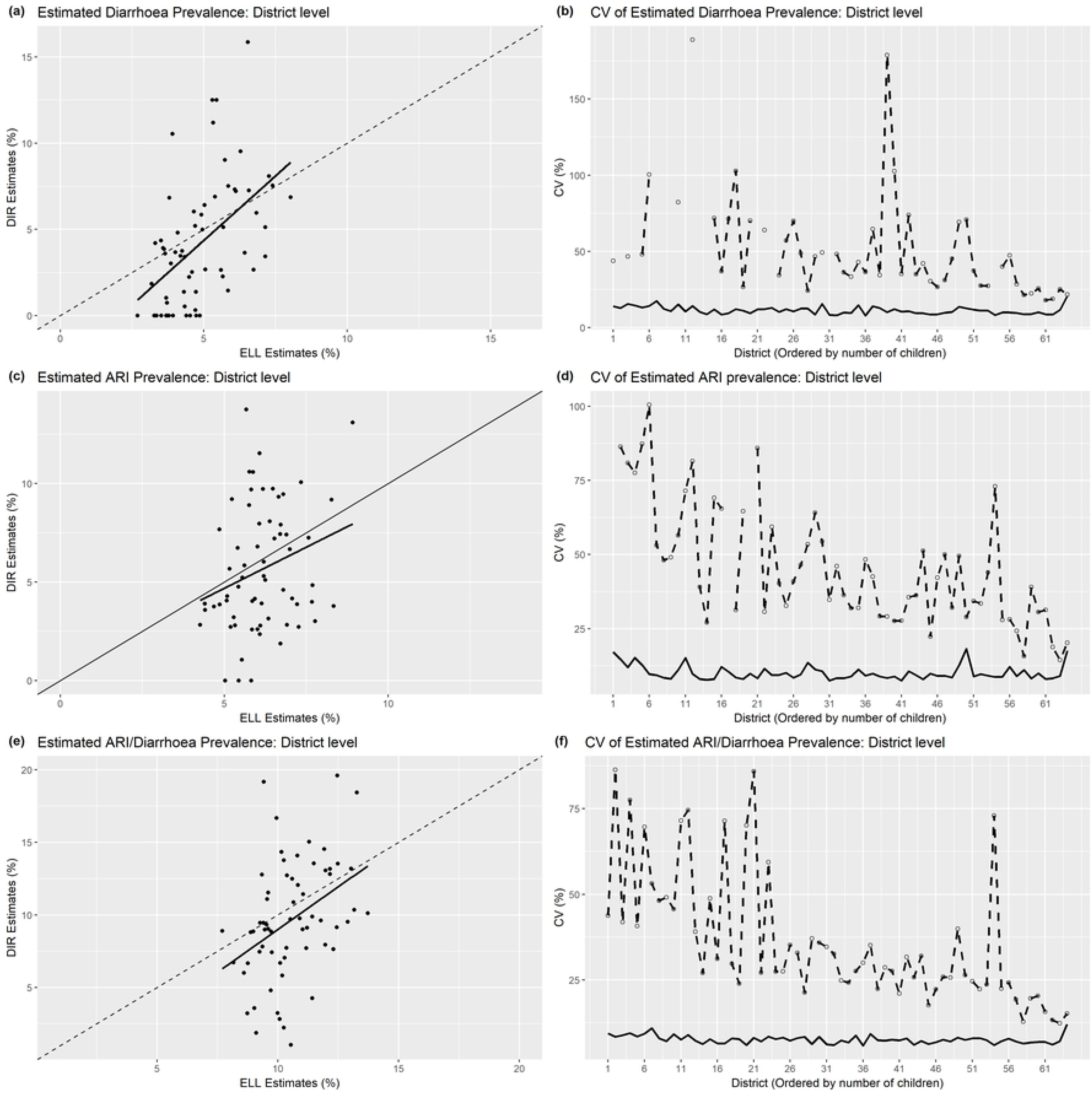
Bias diagnostic plot with y=x line (dashed) and regression line (solid) for estimated prevalence of diarrhoea, ARI, and ARI/diarrhoea among under-5 children at district level in Bangladesh (plots (a), (c), (e) respectively) by the ELL estimator and the corresponding coefficient of variations (CV) against the district index ordered by the number of total children (plots (b), (d), (f) respectively) estimated by direct (dashed line) and ELL (solid line) estimators

**Table 3.**
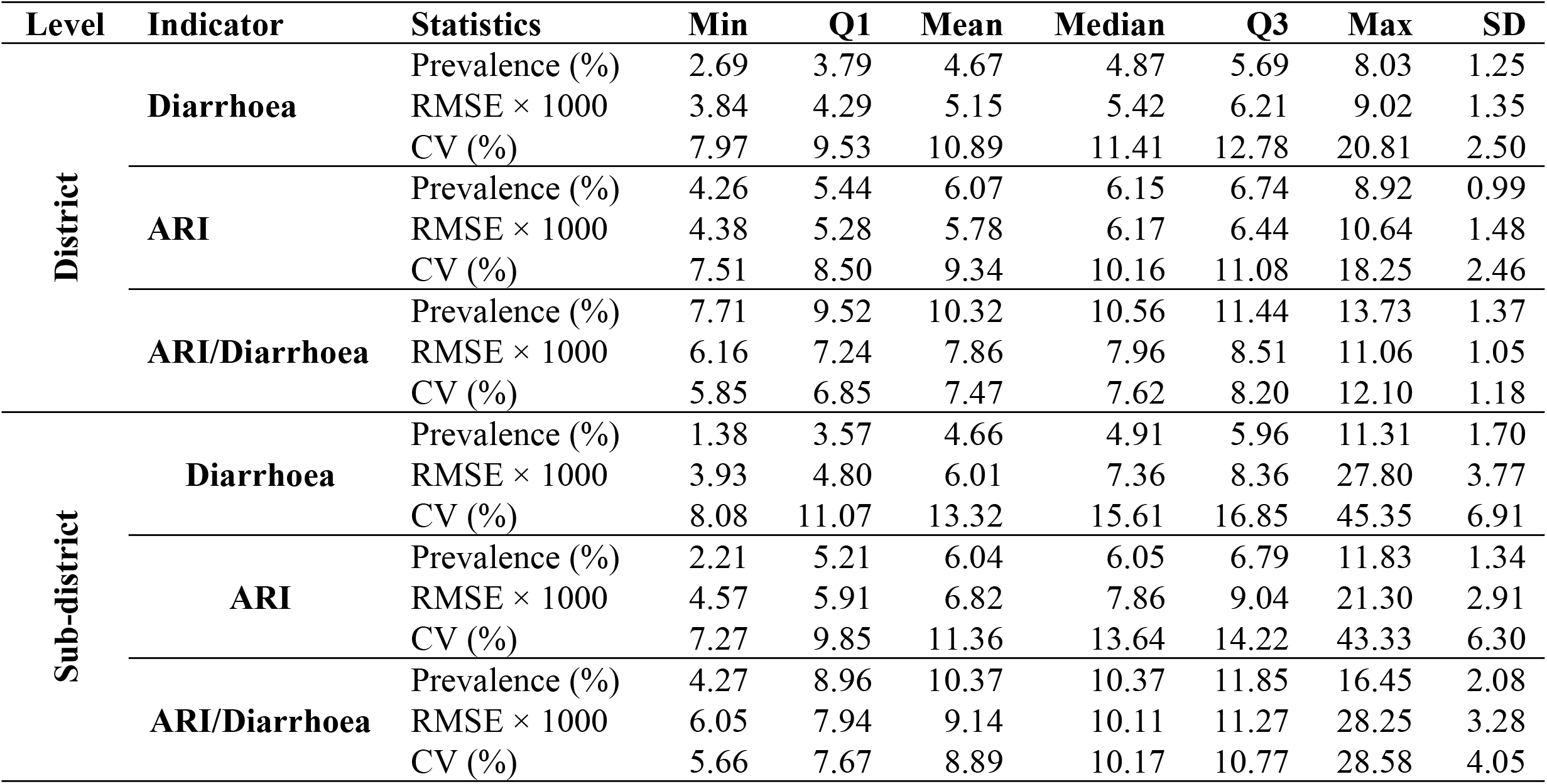
Summary statistics of the estimated diarrhoea, ARI, and ARI/diarrhoea prevalence among under-5 children at district and sub-district level with their estimated root mean squared errors (RMSE) and coefficient of variation (CV%) using ELL estimator.

Since the survey data covers only 396 out of 544 sub-districts and sub-district specific sample sizes are too small, only the ELL estimator is applied to calculate sub-district specific prevalence. Summary statistics of the estimated sub-district level prevalence of diarrhoea, ARI and ARI/diarrhoea shown in Table 3 indicate that the mean prevalence are respectively 4.91% (SD: 1.70%), 6.05% (SD: 1.34%) and 10.37% (SD: 2.08%). Maximum sub-district level prevalence was 11.3% for diarrhoea, 11.8% for ARI and 16.5% for ARI/diarrhoea. It is observed that about 25% sub-districts have more than about 6.0% and 7.0% prevalence of diarrhoea and ARI respectively, while about 75% sub-districts have more than 9.0% prevalence of ARI/diarrhoea. Sub-district level estimates can be considered efficient based on the CV estimates, of which 75% are below 17% for diarrhoea, 14% for ARI and 11% for ARI/diarrhoea.

For identifying the vulnerable hotspots of the considered three health indicators, district and sub-district level maps of Bangladesh are generated using the corresponding estimates calculated by the ELL estimator. In the maps, the seven-shaded colours indicate the distribution of district/sub-district as 0-10%, 10-20%, 20-40%, 40-60%, 60-80%, 80-90%, and 90-100% in ascending order based on the prevalence of the corresponding indictor. Maps in Fig 4 show the spatial distributions of diarrhoea prevalence at district and sub-district levels. District level map shows that the districts of western-south (*Khulna* region) and western (*Rajshahi* region) parts had lower diarrhoea prevalence, while the districts of northern (*Mymensingh* region), north-eastern (*Sylhet* region), south-eastern (*Chittagong* region) and coastal regions of *Barisal* division had comparatively higher diarrhoea prevalence (more than 6%). More specifically, the highest diarrhoea prevalence is found in Cox’s Bazar (8%) and lowest in *Joypurhat* district (3%). Interestingly, “*Chapi Nawabganj*” district lying in the very Western part was highly vulnerable to diarrhoea prevalence, though its neighbouring districts had significantly lower prevalence. The sub-district level map reveals the similar spatial distribution as district level map but it exactly shows the main micro-level hotspots of diarrhoea prevalence. The highly vulnerable sub-districts with prevalence of 7.3-11.4% are mostly in the *Sylhet* and *Chittagong* regions, which are highly prone to floods every year. However, it is observed that there were some sub-districts with higher diarrhoea prevalence belong to the districts with lower diarrhoea prevalence in the district-level map (as for example two sub-districts in *Chuadanga* and *Jhenaidah* districts). Also, the sub-district level map indicates that only two sub-districts of *Chapai Nawabganj* district are main hotspots for its higher diarrhoea prevalence.

**Fig 4:**
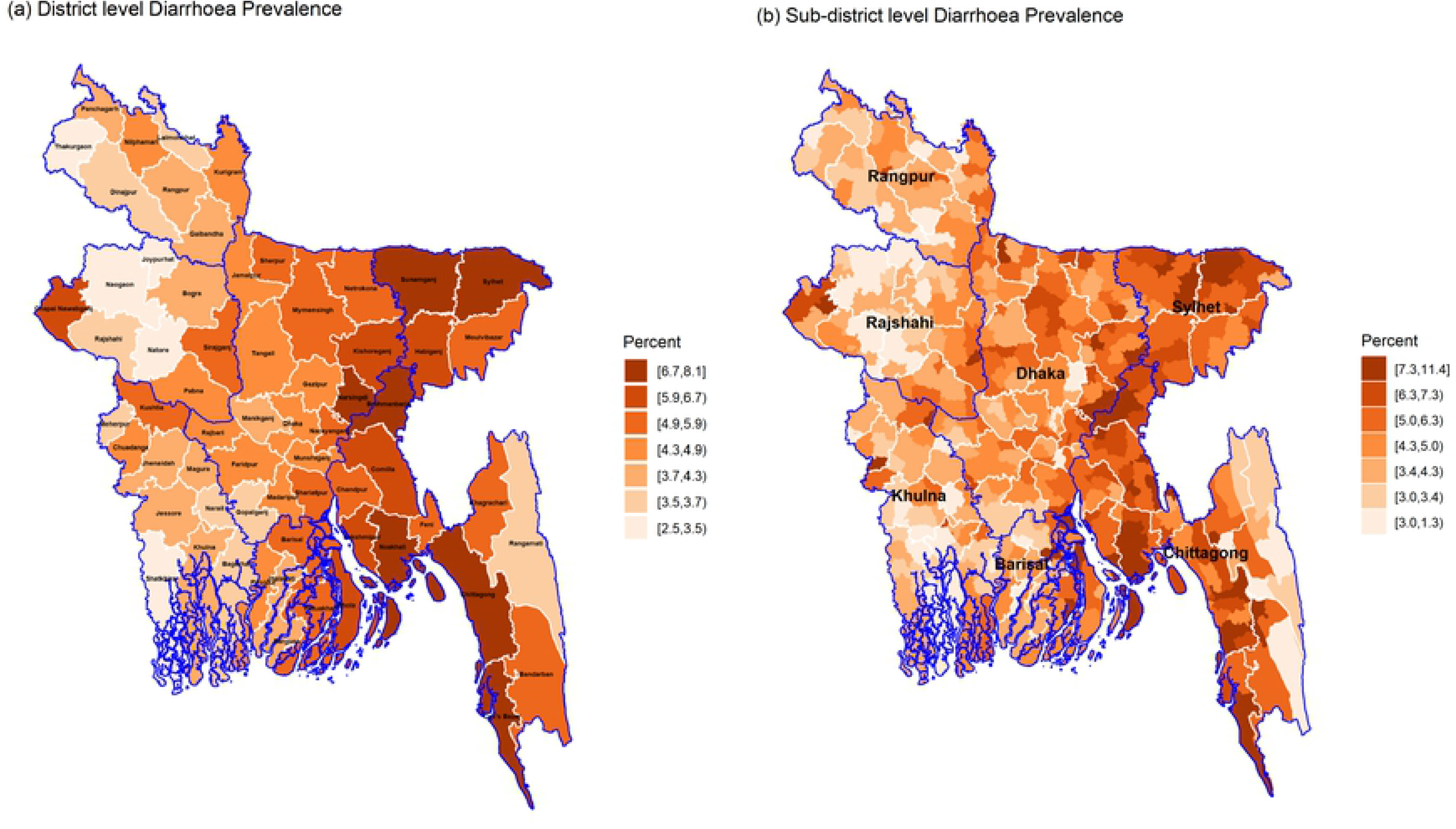
District and sub-district level hotspots of diarrhoea prevalence among under-5 children of Bangladesh (blue and white borderlines refer respectively to division and district areas while no border for sub-districts)

The district and sub-district level prevalence of ARI are mapped in Fig 5. The district level map identifies *Chandpur*, *Comilla*, *Feni*, *Noakhali* and *Lakshmipur* districts of *Chittagong* division, *Khulna* district of Khuna division and Jhalokathi district of *Barisal* division as the highly vulnerable districts for ARI prevalence (7.5-9.0%). While the sub-district level map shows the hotspots with 7.6-11.9% ARI prevalence not only belonged in the above-mentioned districts but also with other districts lying mostly in the whole costal southern part of Bangladesh. Also, the children of Hill Tracts area of *Chittagong* division (mainly *Khagrachari* and *Rangamati* districts) are highly vulnerable to ARI prevalence than to diarrhoea prevalence. Both district and sub-district level maps show hotspots with ARI prevalence of 6.0-7.5% are scattered all over the country except the central region of country.

**Fig 5:**
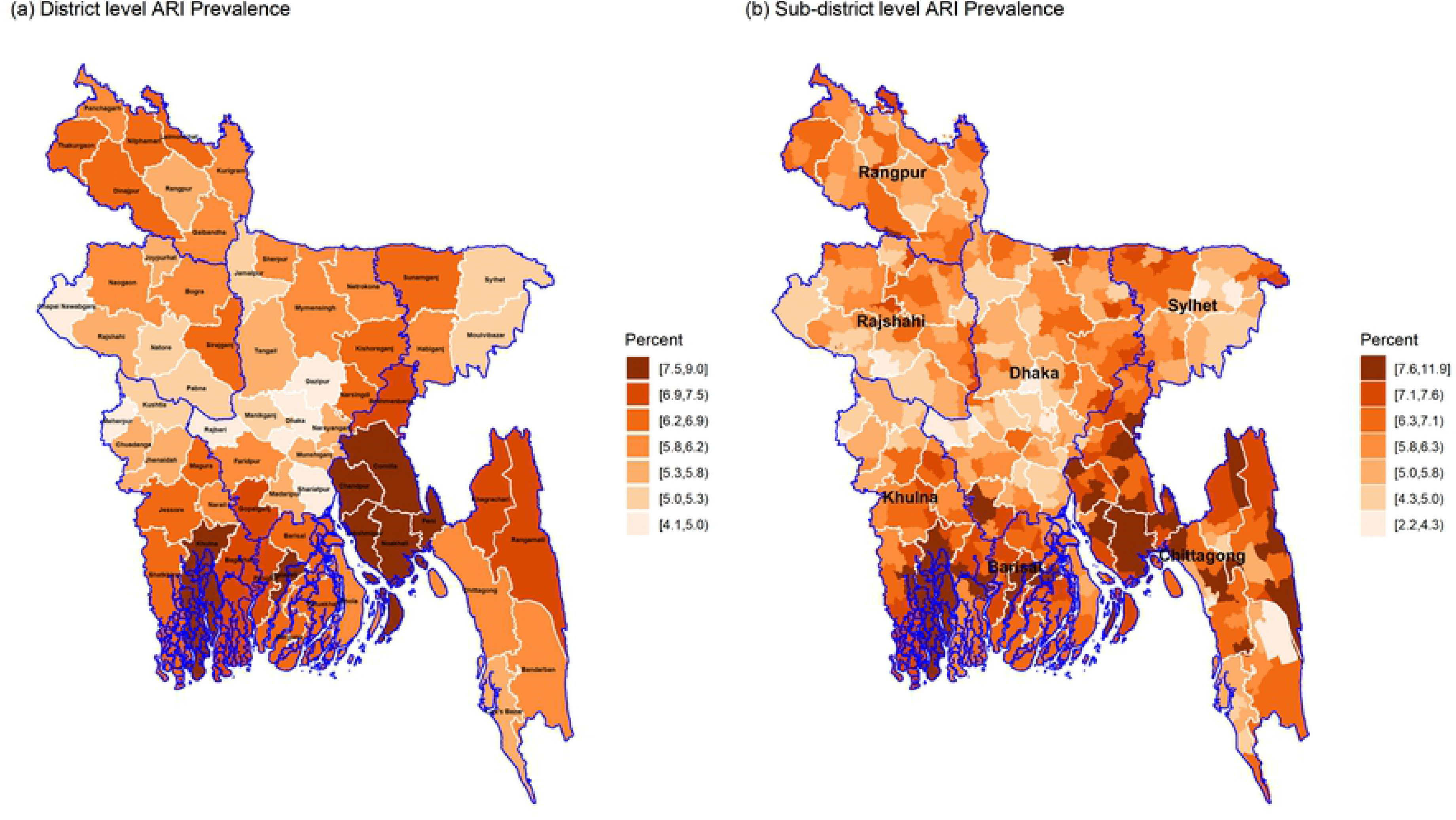
District and sub-district level hotspots of ARI prevalence among under-5 children of Bangladesh (blue and white borderlines refer respectively division and district areas while no border for sub-districts)

The district level distribution of ARI/Diarrhoea prevalence shown in Fig 6 reveals very similar distribution as for the diarrhoea prevalence. Consequently, there are some tendency that the sub-districts vulnerable from diarrhoea were also vulnerable from ARI/diarrhoea (such as *Halishahar*, *Teknaf*, *Companiganj*, *Ramu*, and *Ukhia* sub-districts; please see supplementary S1 File). The sub-district level map indicates that a significant number of sub-districts particularly from *Sunamganj*, *Sylhet*, *Brahmanbaria*, *Narshingdi*, *Noakhali*, *Chittagong*, *Cox’s Bazar*, *Potuakhali* and *Bhola* districts have more than 13% ARI/diarrhoea prevalence. Overall, the highly vulnerable districts/sub-districts are distributed in the right half of the country except the hill tracts districts/sub-districts of *Chittagong* region. The children of *Chapai Nawabganj* district along with its two sub-districts (exceptional in the eastern part of the country) are also found suffering from both diarrhoea and ARI/diarrhoea but not from ARI separately.

**Fig 6:**
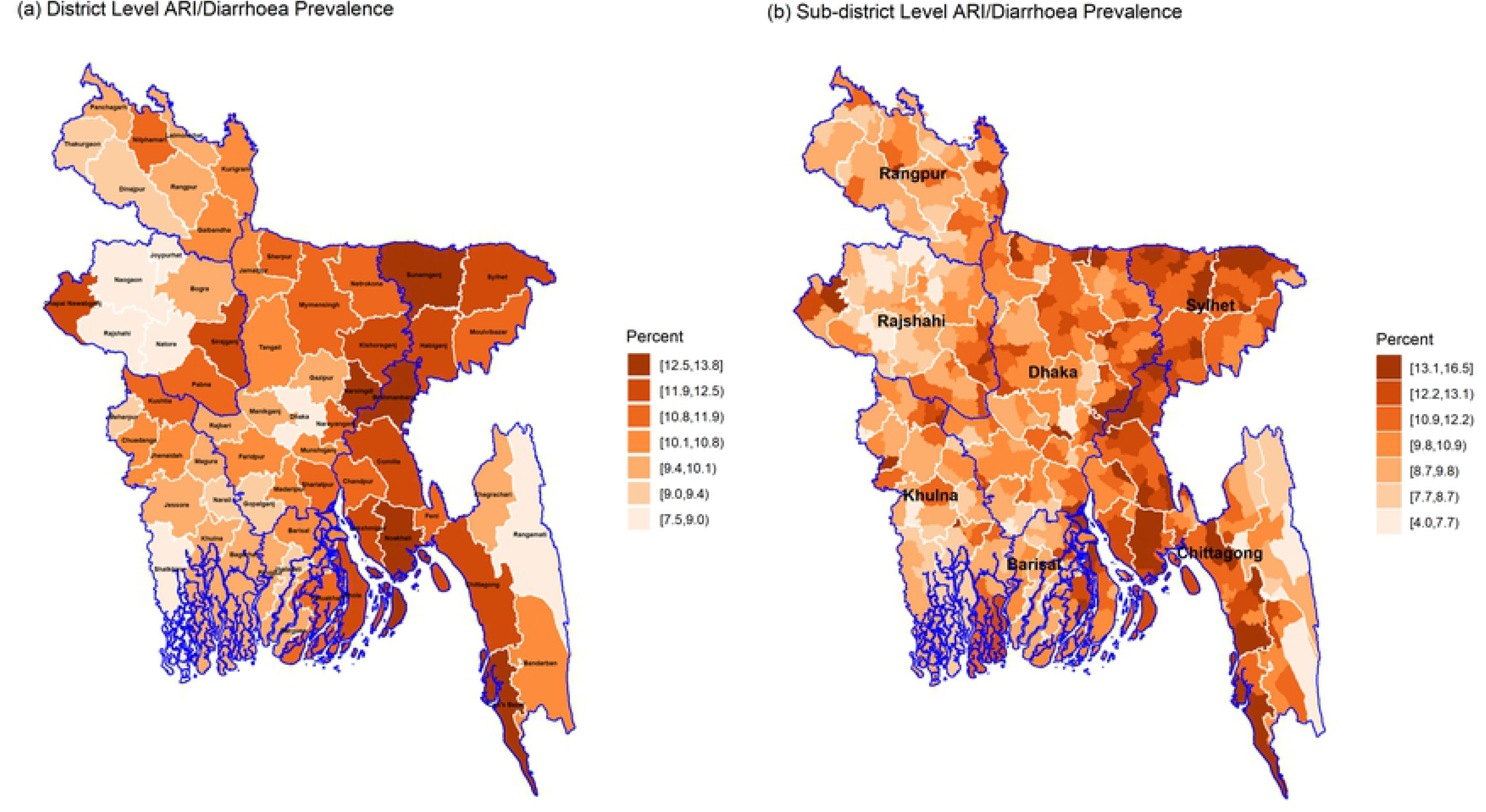
District and sub-district level hotspots of ARI/diarrhoea prevalence among under-5 children of Bangladesh (blue and white borderlines refer respectively division and district areas while no border for sub-districts)

The estimated prevalence of diarrhoea, ARI, and ARI/diarrhoea among under-5 children at district (Zila) and sub-district levels with their 95% CI in supplementary files S1 File and S2 File respectively.

The characteristics of the ELL estimator for binary response variable have been examined by plotting the estimated prevalence and their estimated RMSEs against the sub-district wise total number of children in Fig 7 (plot (a) and (b)). For diarrhoea and ARI/diarrhoea indicators, the estimated prevalence had tendency to increase exponential with the number of under-5 children, while the RMSEs are found to have exponential decreasing trend but remains stable for larger sub-districts. For ARI, the smooth line of RMSE shows an approximate flat U-shaped curve with the number of children, however no specific pattern is observed for the estimated prevalence of ARI against the size of the population. The estimated RMSEs are also plotted against the estimated prevalence to examine the ELL estimator (plot (c) in Fig 7). For diarrhoea, the estimated RMSEs increase exponentially with the prevalence, while the smooth lines of ARI and ARI/diarrhoea show approximately U-shaped pattern. The U-pattern suggest that the RMSEs decrease upto a prevalence level (near about 5% and 9% for ARI and ARI/diarrhoea respectively) and again started to increase after that prevalence level. The distribution of prevalence and their relationship with number of under-5 children may be one of the reasons of this U-pattern graph.

**Fig 7:**
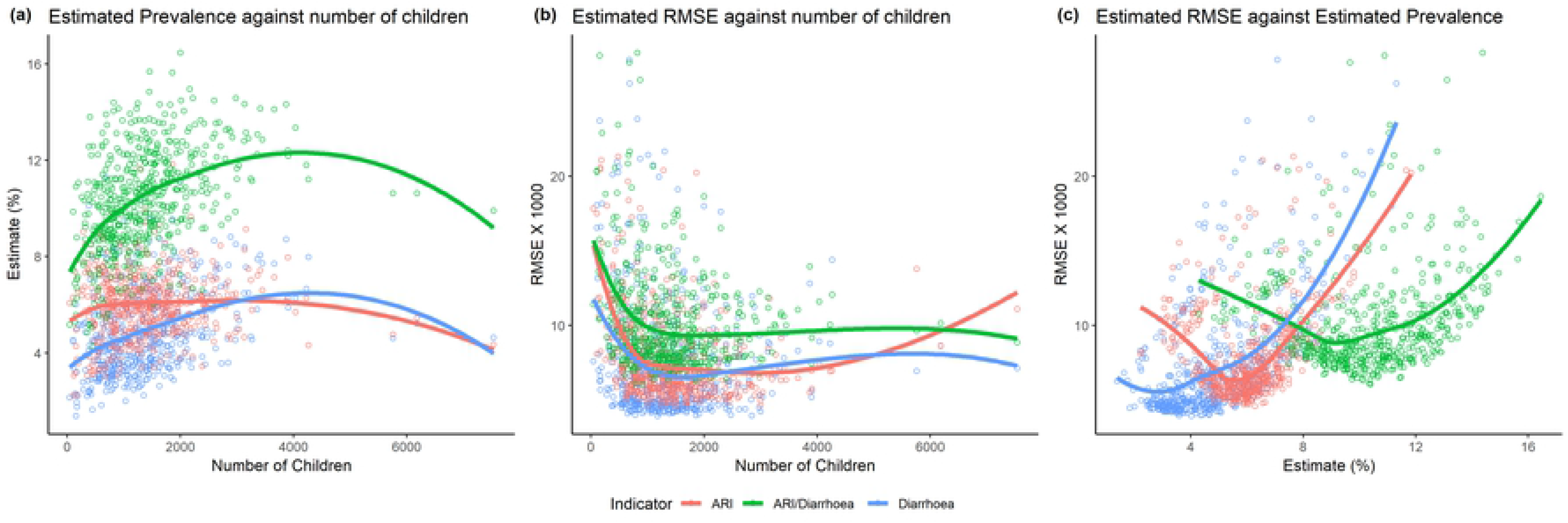
Relationship among the estimated prevalence of diarrhoea, ARI and ARI/diarrhoea among under-5 children at sub-district level, their standard errors (RMSEX1000), and the sub-district specific population size (under-5 children)

Kernel densities of the sub-district specific prevalence of diarrhoea, ARI and ARI/diarrhoea prevalence are plotted in Fig 8. The distribution of diarrhoea prevalence show slightly positive skewness, while the distribution of ARI prevalence shows approximately symmetric but a shape of t-distribution with some extreme prevalence on the right tail. On the other hand, the distribution of ARI/diarrhoea shows slightly negative skewness with very flat tail. These skewed distributions indicate that some sub-districts have unusually high prevalence of diarrhoea (more than 8%) and ARI (more than 10%), also some districts have unusually lower prevalence of ARI/diarrhoea (less than 5%). Hasslet *et al.* [20] found such positively skewed distribution in the SAE study of diarrhoea prevalence in Nepal.

**Fig 8:**
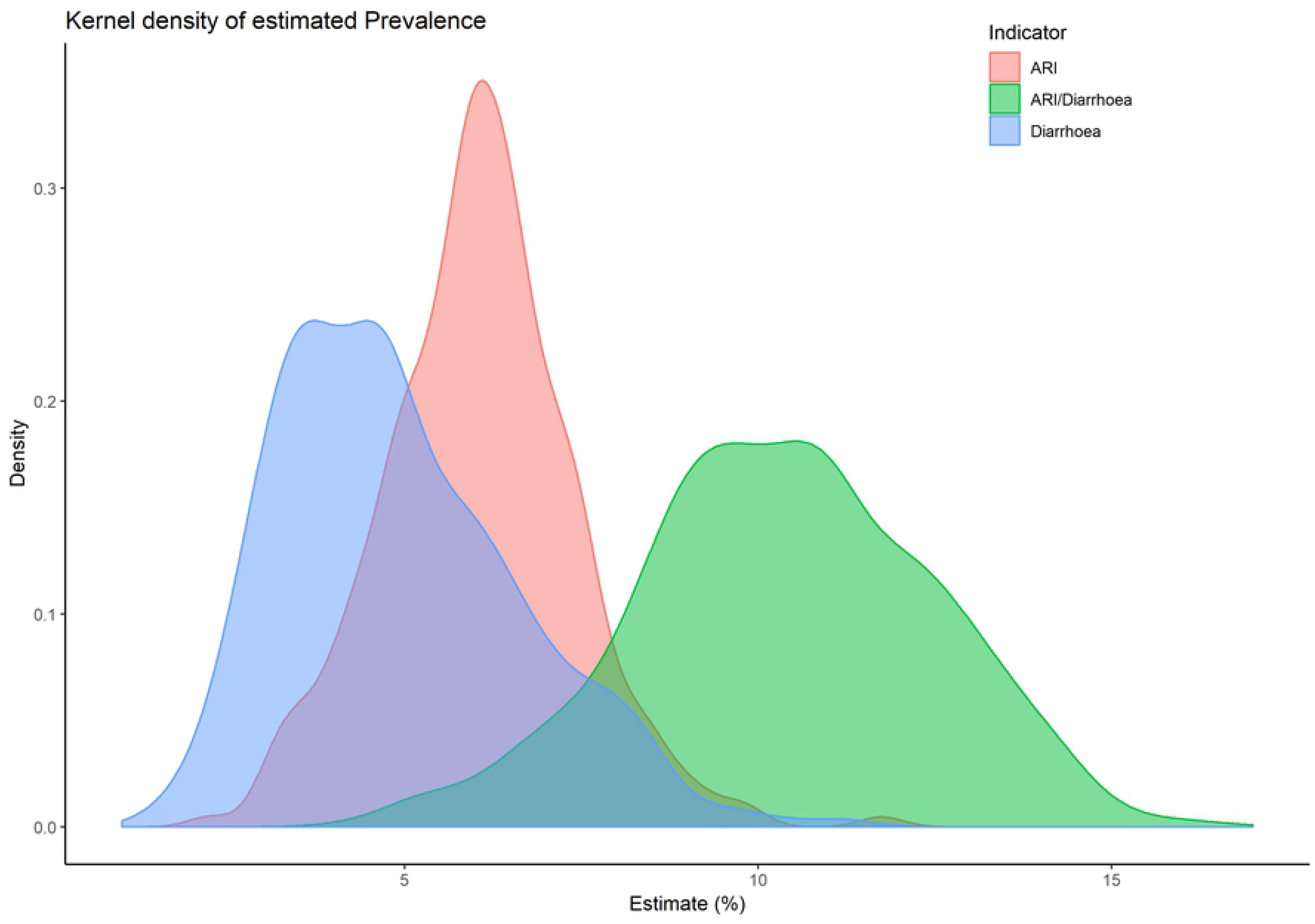
Distribution of sub-district level diarrhoea, ARI, and ARI/diarrhoea prevalence in Bangladesh estimated by the ELL.B estimator

## Conclusions

Diarrhoea and ARI (in the form of pneumonia) are still two leading causes of child deaths not only in Bangladesh but also globally. In Bangladesh, 15% and 6% of 119,000 total under-5 children deaths are still due to diarrhoea and pneumonia diseases in 2015 [26]. Delay in seeking appropriate care and lack of access to multiple sources for treatment are recognized as the underlying risk factors for children death due to diarrhoea and pneumonia in Bangladesh in several research works [12, 13]. Thus, government should focus on the effective interventions of proper life-saving treatment for the children with suspected pneumonia and diarrhoea to achieve the GDDP goals on reducing child mortality due to ARI and diarrhoea. In this respect, identifying the hotspots of children diarrhoea and ARI prevalence at the lower administrative units would be helpful for the government. Taking proper initiative for providing the proper life-saving treatments in those hotspots may reduce the mortality due to these two leading causes.

This study aims to identify the hotspots of diarrhoea and ARI prevalence at two disaggregate levels in Bangladesh through application of a SAE method for dichotomous response variable. The application provides district and sub-district levels prevalence of ARI, diarrheal, and ARI/diarrhoea episodes with their accuracy measures. The study findings confirmed that though the national level and division level estimates of the three health indicators seem very low, there are significant inequalities among the districts and sub-districts. The inequality increases when the aggregation level goes down to sub-district level from division level.

The comparison of the interactive maps at district level suggests that children living in the southern and north-eastern parts of Bangladesh have higher tendency to experience with the incidence of diarrhoea and ARI related diseases compared to those children living in the central, western and north-western parts. District level maps of the three indicators indicate that children of five districts (*Sunamganj*, *Brahmanbaria*, *Narshindi*, *Noakhali* and *Cox’s Bazar*) are highly vulnerable to diarrhoea as well as ARI/diarrhoea, while only those of *Noakhali* district are highly susceptible to all the three indicators. The sub-district level maps suggest that the higher risk of occurring child morbidity was not restricted only to those sub-districts belonging to the highly vulnerable districts but also in some districts with less vulnerability. For ARI, moderate to highly vulnerable sub-districts are found over all regions except the sub-districts very close to the capital city *Dhaka*.

A comparison between district and sub-district level maps identifies those sub-districts for which the prevalence of a specific district becomes higher (as for example, three sub-districts of *Chapai Nawabganj* have diarrhoea prevalence of more than 7%, while the other sub-districts have prevalence below 4%). Das, Chandra and Saha [22] also found that *Chapai Nawabganj* had significantly higher diarrhoea prevalence but it was not possible to say for which sub-districts. Sub-district level prevalence of this study indicate that children of “*Gomastapur*”, “*Nawabganj Sadar*” and “*Shibganj*” are more vulnerable to diarrhoea prevalence. It is also observed that some districts with very low prevalence of an indicator (say, diarrhoea) have sub-districts with considerably higher prevalence (for example, diarrhoea prevalence for *Rangamati* district is only 3.7% but it has a sub-district with prevalence of 5.2%).

The findings suggest that the ELL estimators for the three dichotomous response variables are providing unbiased and consistent estimates compared to the direct estimator at both district and sub-district level. The benefits of this ELL estimator is that an appropriate multilevel model developed at the most detailed model can be used for estimating target parameters at the several aggregation levels. In this study, a suitable multilevel model for each of the health indicators has been developed using the survey data and some contextual variables extracted from the 5% census data, and then the prediction has been done using the children level explanatory variables available in the Census data. If full census data can be used the estimator will be more consistent at sub-district level and even the indicators can be estimated at lower administrative units like *Union* and *Mauza* levels.

As limitations of this study, two issues concerned the authors: normality issues of the cluster-specific residuals and the shrinkage of the estimated target parameters to the average. The shrinkage can be removed if some contextual variables at district and sub-district level can be incorporated in the final model. The authors tried to improve the model by incorporating such variables, but the incorporated contextual variables are not found significant in the final model. In further study, the ELL estimator used in this study can be compared to the estimator based on the model developed at the target aggregation level such as district or sub-district level. For sub-district level, the problem is that about 40% sub-districts are not available in the survey data and so sub-district level residuals cannot be used for prediction for all sub-districts as required in the empirical Bayes estimator described in Molina and Rao [27] utilizing a GLMM model. Ultimately, a synthetic type estimator like ELL will be required for those non-sampled sub-districts.

Children malnutrition status is highly correlated with their recent experience with the occurrence of diarrhoea and ARI related diseases. Evidence from several child malnutrition studies [28, 29] suggest that the children recently suffered from diarrhoea and ARI are more likely to be either wasted (indicator of acute malnutrition) or underweight (combination of acute and chronic malnutrition). The national target of reducing wasting at 5% by 2025 [30] will be achievable if the prevalence of ARI and diarrhoea can be reduced as well. Haslett, Jones and Isidro [31] conducted a small area study on child undernutrition (underweight: lower weight compared to age) in Bangladesh using the Child and Mother Nutrition Survey of Bangladesh 2012 and full data of Census 2011. They found that the children living in the north-eastern (*Sylhet* and *Mymensingh* regions) and the south-eastern (costal parts of *Chittagong* region) parts of Bangladesh are vulnerable to both underweight and severely underweight (Apendix D.3 of Haslett, Jones and Isidro [31]). These spatial distributions of underweight and severe underweight are seemed comparable to the spatial distribution of ARI/diarrhoea generated in this study (map (b) in Fig 6). The comparison of these spatial distributions reveals the inter-relationship between child undernutrition and incidence of recent diarrhoea and ARI. Thus, combination of this small area study on child diarrhoea and ARI prevalence with the study of child undernutrition [31] might guide the stakeholders for taking proper initiatives at the lower administrative levels for reducing the vulnerability of child morbidity and undernutrition simultaneously. The interventions at the lower administrative units might be followed up so that the children in risk of severe diarrhoea and pneumonia can get the access of life-saving treatment timely, which save the children as well as reduce the rate of child mortality due to diarrhoea and ARI.

## Funding details

Research work was supported by SUST Research Center, Shahjalal University of Science & Technology for the period of 2017-2018.

## Disclosure statement

Authors do not have any conflict of interest.

## Acknowledgment

The authors would like to acknowledge the valuable comments and suggestions of the Professor Emeritus Stephen Haslett, Professor of Statistics, Institute of Fundamental Science & Centre for Public Health Research, Massey University. His suggestions led to a considerable improvement in the paper.

## Supporting information

**S1 File 1: Estimated prevalence of diarrhoea, ARI, and ARI/diarrhoea among under-5 children at district (Zila) level with their 95% CI**

**S2 File 1: Estimated prevalence of diarrhoea, ARI, and ARI/diarrhoea among under-5 children at sub-district (Up-zila) level with their 95% CI**

**Appendix Table 1.** Regression Coefficients (p-values) of the fitted model for diarrhoea, ARI and ARI/diarrhoea based on BDHS 2011 and Census 2011 data.

| Explanatory Variables | Diarrhoea |  | ARI |  | ARI/Diarrhoea |  |
| --- | --- | --- | --- | --- | --- | --- |
| | $\beta$ | p-value | $\beta$ | p-value | $\beta$ | p-value |
| (Intercept) | -3.408 | 0.002 | -2.158 | 0.000 | -2.827 | 0.000 |
| Age 1 | 0.164 | 0.330 |  |  |  |  |
| Age 2 | -0.523 | 0.009 |  |  |  |  |
| Age 3 | -0.498 | 0.007 |  |  |  |  |
| Age 4 | -0.832 | 0.000 |  |  |  |  |
| pliterate |  |  |  |  | -0.435 | 0.018 |
| pkids714 |  |  | -0.808 | 0.021 |  |  |
| Urban | 0.837 | 0.081 |  |  |  |  |
| Urban:hfem | 1.342 | 0.033 |  |  |  |  |
| Urban:pliterate | -1.192 | 0.014 |  |  |  |  |
| Male: AGE 0 |  |  | 0.877 | 0.000 | 0.826 | 0.000 |
| Female: AGE 0 |  |  | 0.434 | 0.081 | 0.440 | 0.017 |
| Male: AGE 1 |  |  | 0.918 | 0.000 | 0.945 | 0.000 |
| Female: AGE 1 |  |  | 0.602 | 0.013 | 0.685 | 0.000 |
| Male: AGE 2 |  |  | 0.585 | 0.018 | 0.519 | 0.005 |
| Female: AGE 2 |  |  | 0.596 | 0.014 | 0.421 | 0.023 |
| Male: AGE 3 |  |  | 0.456 | 0.057 | 0.324 | 0.072 |
| Female: AGE 3 |  |  | 0.297 | 0.239 | 0.123 | 0.522 |
| Male: AGE 4 |  |  | 0.494 | 0.042 | 0.143 | 0.451 |
| Urban: Age 1 | 0.263 | 0.441 |  |  |  |  |
| Urban: Age 2 | 0.758 | 0.039 |  |  |  |  |
| Urban: Age 3 | -0.076 | 0.854 |  |  |  |  |
| Urban: Age 4 | 0.451 | 0.270 |  |  |  |  |
| Urban: *Barisal* |  |  | -0.167 | 0.580 | 0.150 | 0.497 |
| Urban: *Chittagong* | -0.490 | 0.167 | -0.425 | 0.080 | -0.218 | 0.213 |
| Urban: *Dhaka* | -1.127 | 0.005 | -0.299 | 0.325 | -0.691 | 0.001 |
| Urban: *Khulna* | -0.826 | 0.066 | 0.115 | 0.656 | -0.023 | 0.912 |
| Urban: *Rajshahi* | -0.902 | 0.047 | 0.074 | 0.790 | -0.199 | 0.370 |
| Urban: *Rangpur* | -0.958 | 0.044 | 0.088 | 0.749 | -0.085 | 0.704 |
| Urban: *Sylhet* | -0.557 | 0.155 | -1.026 | 0.008 | -0.502 | 0.027 |
| **Sub-district Level Contextual Variables** |  |  |  |  |  |  |
| pemp15_subdist |  |  | -2.865 | 0.043 |  |  |
| plit_subdist | -2.832 | 0.000 | -9.018 | 0.005 | -1.239 | 0.025 |
| phhlit_subdist |  |  | 6.056 | 0.003 |  |  |
| phhcssc_subdist |  |  | 3.985 | 0.081 |  |  |
| pempag_subdist | -5.112 | 0.000 |  |  |  |  |
| pfcmced_subdist | 2.628 | 0.005 |  |  | 1.440 | 0.027 |
| padnowork_subdist |  |  |  |  | -3.316 | 0.000 |

**Appendix Table 2.**
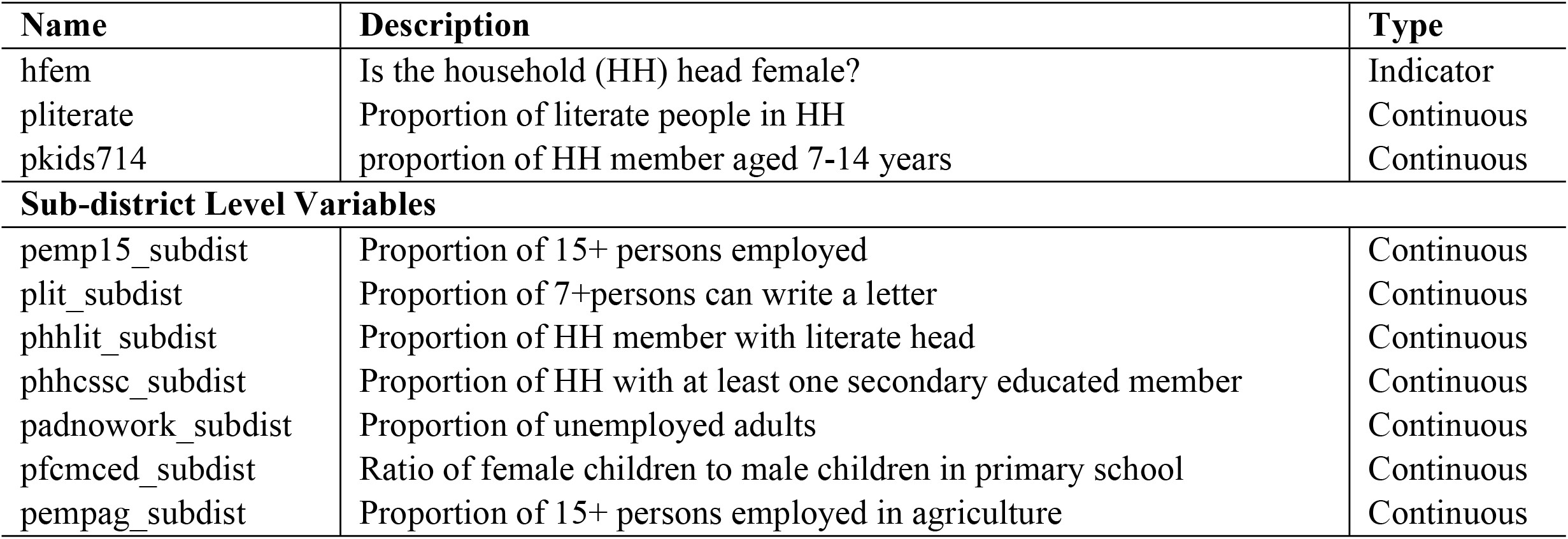
Description of the contextual variables at different levels calculated from BDHS 2011 and Census 2011 data.

